# OxoScan-MS: Oxonium ion scanning mass spectrometry facilitates plasma glycoproteomics in large scale

**DOI:** 10.1101/2022.06.01.494393

**Authors:** Matthew E. H. White, D. Marc Jones, Joost de Folter, Simran Kaur Aulakh, Helen R. Flynn, Lynn Krüger, Vadim Demichev, Pinkus Tober-Lau, Florian Kurth, Michael Mülleder, Véronique Blanchard, Christoph B. Messner, Markus Ralser

## Abstract

Protein glycosylation is a complex and heterogeneous post-translational modification. Specifically, the human plasma proteome is rich in glycoproteins, and as protein glycosylation is frequently dysregulated in disease, glycoproteomics is considered an underexplored resource for biomarker discovery. Here, we present OxoScan-MS, a data-independent mass spectrometric acquisition technology and data analysis software that facilitates sensitive, fast, and cost-effective glycoproteome profiling of plasma and serum samples in large cohort studies. OxoScan-MS quantifies glycosylated peptide features by exploiting a scanning quadrupole to assign precursors to oxonium ions, glycopeptide-specific fragments. OxoScan-MS reaches a high level of sensitivity and selectivity in untargeted glycopeptide profiling, such that it can be efficiently used with fast microflow chromatography without a need for experimental enrichment of glycopeptides from neat plasma. We apply OxoScan-MS to profile the plasma glycoproteomic in an inpatient cohort hospitalised due to severe COVID-19, and obtain precise quantities for 1,002 glycopeptide features. We reveal that severe COVID-19 induces differential glycosylation in disease-relevant plasma glycoproteins, including IgA, fibrinogen and alpha-1-antitrypsin. Thus, with OxoScan-MS we present a strategy for quantitatively mapping glycoproteomes that scales to hundreds and thousands of samples, and report glycoproteomic changes in severe COVID-19.

## Introduction

The proteomes of liquid biopsies and peripheral body fluids, in particular blood plasma or serum, are an emerging source of biomarkers, bearing potential for novel diagnostic, prognostic, and predictive applications.^1,2^ The plasma proteome contains important nutrient response proteins and components of the immune system, whose concentration and activity reflect the physiological condition of the individual and which are therefore important for precision medicine.^3^ Technologies facilitating the quantification of the plasma proteome in large sample series, using mass spectrometry2 or with the affinity-reagent based Olink^4^ and SomaScan^5^ platforms, have opened exciting new avenues to better link genetic diversity and disease phenotypes at the epidemiological scale.^6^ However, the activity and function of proteins depends not only on their abundance, but also post-translational modifications, protein-protein and protein-small molecule interactions, processes that themselves depend on whether a protein is modified.^7^ As a consequence, abundance measurements alone capture only part of the human physiology represented by the plasma proteome, creating a need to develop methods that can address post-translational modifications and proteoforms at cohort scale.

An important reservoir for biomarker discovery is glycoproteomics. Protein glycosylation is abundant and diverse in plasma, and altered glycosylation has been observed in response to a variety of disease states, for example, prostate-specific antigen in prostate cancer and alpha-1-acid glycoprotein in sepsis.^8–10^ Therefore, there is an increasing demands for approaches that allow the sensitive and quantitative profiling of blood plasma, where the high abundance and diversity of protein glycosylation plays a vital role in regulating the structure and function of both soluble and cell-surface proteins.^11^ Liquid chromatography-mass spectrometry-based (LC-MS) proteomic technologies are widely applied in the identification and quantification of post-translational modifications in cell- and tissue-derived samples.^7,12–17^ Furthermore, through advances in sample preparation and novel data acquisition strategies, MS-based technologies have also reached a level of robustness and throughput for large-scale, high-throughput investigations that involve the measurement of thousands of samples.^18–22^

The study of intact glycopeptides at scale still presents a number of analytical challenges, however. A large proportion of glycoproteins have multiple glycosylation sites (macroheterogeneity), at each of which there is a large range of possible glycan structures (microheterogeneity). The abundance of a given (glyco)protein is therefore comprised of various individual glycoforms at lower respective concentrations, necessitating a highly sensitive analytical approach.^23,24^ Furthermore, co-elution of unmodified peptides reduces sensitivity via ion suppression, and for data-dependent acquisition, by reducing the time spent by the instrument specifically sampling glycopeptides.^25^ These effects are compounded by the poorer ionisation efficiency of glycopeptides relative to their unmodified counterparts.^26^ A number of glycoprotein/glycopeptide enrichment strategies have been developed to minimise the challenges of intact glycopeptide analysis.^27^ These reach excellent depth on individual samples, but increased cost, handling time and potential batch effects limit their application on large cohort studies. Data-independent acquisition (DIA) methods, such as SWATH-MS, have been increasingly applied in the analysis of large proteomic sample series.^28–31^ In glycoproteomics, DIA approaches have been applied to assess glycosite occupancy of enzymatically deglycosylated peptides^32–35^, and more recently, facilitated the post-acquisition analysis of intact glycopeptides, either by targeted extraction of abundant Y-type (intact peptide with glycan fragments of various sizes) ions^36–40^ or by searching against spectral libraries.^15,41–43^ These approaches yield remarkable depth in comparative analyses and in generating spectral libraries, generally using higher-collisional dissociation (HCD) and/or electron-based fragmentation techniques.^43–45^ Precursor ion scanning and DIA methods have further been applied to quantify oxonium ions, small, singly-charged fragment ions ubiquitously found in glycopeptide HCD/CID MS/MS spectra^46–48^ in biotherapeutics and purified glycoproteins, as well as complex biofluids.^36,39,49–54^

We herein focused on building on these technological advances for developing a glycoproteomic screening platform for high-throughput studies. In order to reach throughput at low cost, we take a two-step approach that separates glycopeptide quantification from sequence assignment. Specifically, in a screening step, we exploit the sensitive detection and quantification of diagnostic oxonium ions for individual glycopeptide features and combine it with an extra scanning quadrupole dimension, as introduced with Scanning SWATH18, to assign precursors to quantified oxonium ions. The information obtained from the scanning dimension facilitates the matching of precursor and MS/MS information between OxoScan- and DDA-glycoproteomics data for identification of the glycopeptides in the second step. In combining the scanning approach for screening and quantifying oxonium ions with a subsequent targeted approach for identification, we reach high levels of sensitivity, acquisition speed and quantitative performance in the detection of glycopeptide features.

We demonstrate the application of OxoScan-MS using micro-flow chromatography by identification of 30 IgG glycoforms without predefined compositional knowledge, and further validate glycopeptide signal specificity and quantitative performance in tryptic digests of human plasma and serum. Moreover, we applied OxoScan-MS to generate a plasma glycoproteome for a cohort of 30 hospitalised COVID-19 patients and 15 healthy controls, in technical triplicates. On the clinical samples (neat citrate plasma), our approach quantified >1,000 glycopeptide features in just 19 minutes of active chromatographic separation across 164 samples, measured in just three days of instrument time. We selected a subset of quantitatively interesting glycopeptide features as potential glyco-biomarkers from the COVID-19 cohort and utilised an orthogonal acquisition approach (DDA HCD-pd-ETD) to perform glycopeptide identification. Critically, our platform captures quantitative biological variation in a plasma cohort. Follow-up analysis of glycopeptide features-of-interest and integration with protein-level data identified potential biomarkers and differential glycan regulation with increasing COVID-19 disease severity. Thus, OxoScan-MS facilitates glycoproteomics on neat plasma and in large scale, and to our knowledge, we report the first cohort-level plasma glycoproteomic analysis of severe COVID-19.

## Results

### Precursor assignment using a scanning quadrupole allows untargeted glycopeptide profiling in OxoScan-MS

We recently described a DIA-based scanning quadrupole acquisition method, Scanning SWATH, in which a scanning quadrupole (Q1) facilitates assignment of precursor masses by time-dependent fragment ion detection in a DIA-MS experiment.^18^ In OxoScan-MS, the scanning dimension allows the extraction of a ‘Q1 profile’ for fragment ions as the precursor enters and exits the sliding Q1 isolation window, centred on the precursor m/z. We demonstrate that selectively extracting Q1 profiles of oxonium ions, which are produced when glycans fragment under CID/HCD conditions^46–48^, allows detection of glycopeptide precursors, even in the presence of co-eluting, highly abundant, unmodified peptides (Figure 1a,b). By overlaying Q1 traces with MS1 spectra, accurate masses can be assigned (Figure 1c). As extracted ion chromatograms show glycopeptide elution in the chromatographic dimension (Figure 1d), selectively extracting oxonium ion chromatograms across the entire precursor range generates a 2-dimensional matrix of glycopeptide signals, even in complex samples containing mostly unmodified peptides (Figure 1e). Not only does this remove the need for pre-defined knowledge of glycopeptide constituents and the biases associated with an empirical spectral library, it also allows relative quantification between samples.

**Figure 1:**
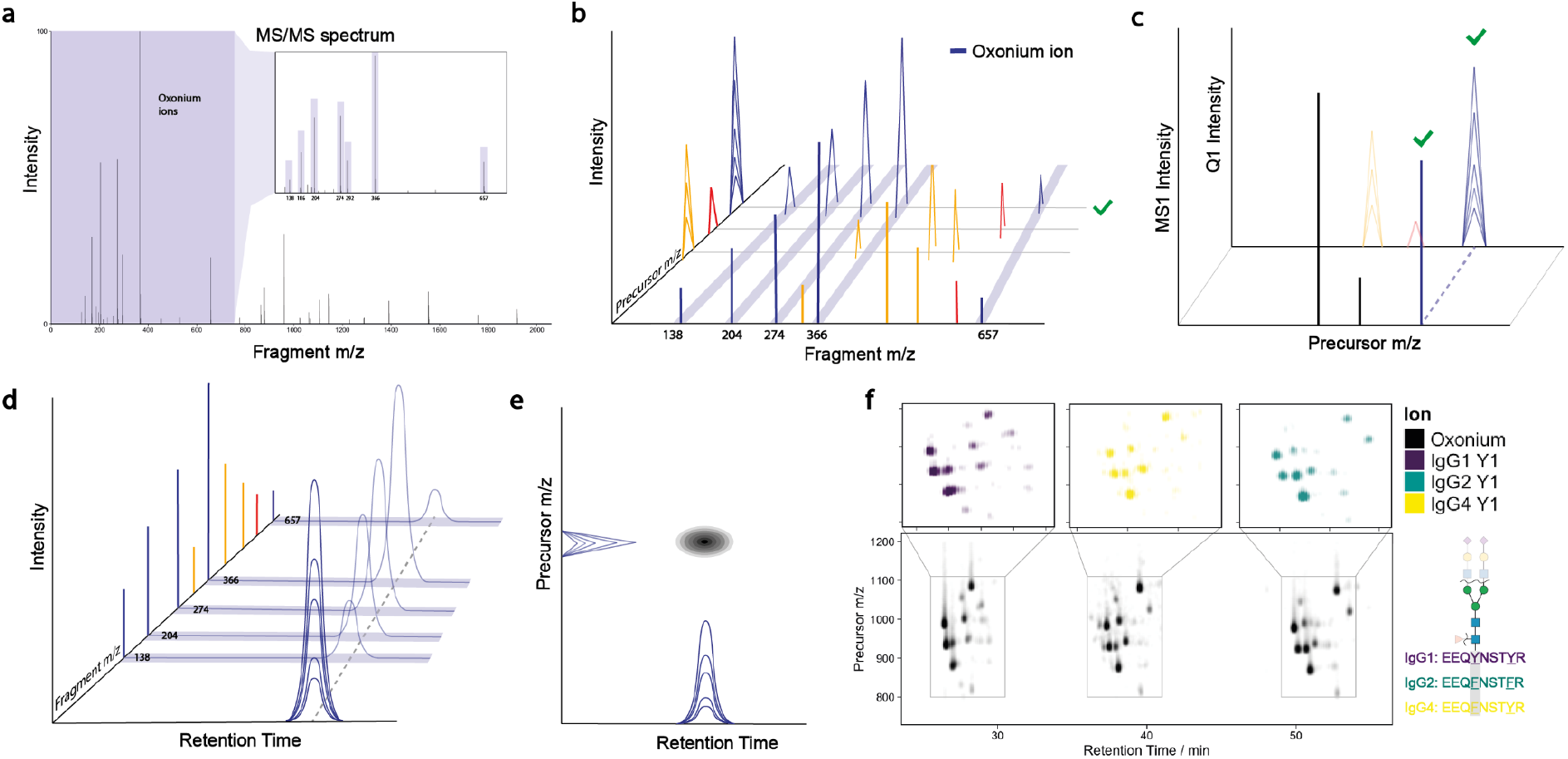
OxoScan-MS exploits a scanning quadrupole for selective glycopeptide profiling by precursor assignment of glycan-specific ions: **a.** Typical tandem mass spectrometry (MS/MS) spectrum from a glycopeptide fragmented under collision-induced dissociation (CID) conditions. The oxonium ions arising from fragmentation of HexNAc (138.05, 186.08, 204.09), Neu5Ac (274.09, 292.10), HexNAc-Hex (366.14) and HexNAc-Hex-Neu5Ac (657.24) ions are highlighted in the inset spectrum. **b.** Time-dependence of the scanning quadrupole gives a ‘Q1 profile’ of each fragment ion entering and exiting the sliding precursor isolation window, which is centred around the precursor mass. **c.** Glycopeptide precursors can be identified by overlaying oxonium ion Q1 profiles on MS1 spectra. Oxonium ion Q1 peaks are glycopeptide-selective, as co-eluting unmodified peptides do not give rise to oxonium ion fragments. Green tick indicates the glycopeptide precursor, localised by oxonium ion signals in Q1 dimension. **d.** Extracted ion chromatograms (XICs) depict elution of a glycopeptide in the chromatographic time dimension by oxonium ion signals. **e.** Each glycopeptide feature, defined as a glycopeptide in a specific charge state, can be localised to a unique retention time-precursor m/z coordinate, with peak height proportional to peak signal. **f.** Oxonium ion map of IgG 1, 2, and 4 glycopeptides with IgG peptide sequences shown for each subclass on right-hand schematic. The lower panel shows the sum of oxonium ion intensities, where each cluster of spots corresponds to the glycopeptides of each IgG subclass at the conserved N-glycan site and each spot is a specific glycopeptide. For ease of interpretation, intensities for respective subclasses have been scaled separately. Inset panels (above) show the Y1 (peptide + GlcNAc) ions extracted and plotted for each IgG subclass, respectively.

To test the validity of this principle, we first profiled IgG subclasses 1, 2 and 4, purified from human blood serum.^55^ By extracting chromatograms of commonly identified oxonium ions across the acquired precursor range, a ‘glycopeptide map’ visually identified >30 features corresponding to the IgG glycopeptides (Figure 1f, Figure S1a). Matching MS1 features to previously reported MS1 signals of glycopeptides (from MALDI-TOF MS^55^) led to the identification of 30 of these glycopeptide features (Table S1). Importantly, we observed well-documented and reproducible retention time shifts for the glycopeptides of each IgG-subclass, recapitulating known behaviour of different glycans with reverse-phase separations (Figure S1b).^56,57^

Recent studies have shown the utility of Y-type fragment ions for quantification and generation of site-specific glycopeptide information in DIA analysis.^36,38,58^ Based on these observations, we developed a rolling collision energy scheme, such that the MS/MS spectra of each glycopeptide feature also contain useful Y-type fragments for targeted re-analysis. Although these spectra can not (yet) be processed with currently available glycoproteomic search engines, we found that highly abundant fragments of peptides with 1—5 attached sugar molecules (the remainder of the glycans being preferentially fragmented over the peptide backbone) allow identification of features from the same peptide. Indeed, we find that Y1 (peptide + GlcNAc) fragments in particular, when calculated *in silico*^36^ and extracted in DIA-NN31, overlay on their respective oxonium ion features, facilitating the distinction of glycopeptides from different IgG subclasses by their respective peptide sequences (Figure 1f, upper panels). This highlights a key advantage of OxoScan-MS; each run acts as a digital archive of the glycoproteome of a sample that can be retrospectively mined. Consequently, OxoScan-MS leverages the advantages of both a precursor ion scan and SWATH-MS in a single run for untargeted quantification of all glycopeptide features above the oxonium ion limit of detection.

### OxoScan-MS quantifies glycopeptide-specific response for >1100 glycopeptide features from neat plasma in short chromatographic gradients

We next chose to test the performance of our method on human plasma. As a large proportion of plasma proteins are glycosylated, we expected to generate significantly more complex data than that obtained from purified IgG.^59^ Analysis of a plasma sample prepared using a semi-automated, high-throughput sample preparation pipeline^20^ with OxoScan-MS (Figure 2a) produced a complex oxonium ion map with hundreds of visible features (Figure 2b, S1c).^20^ To confirm glycopeptide specificity of oxonium ion signals, we treated the sample with a cocktail of glycosidases (Protein Deglycosylation Mix II, New England Biolabs), which enzymatically cleave most glycan classes from proteins, leaving predominantly de- and non-glycosylated peptides. The glycosidase treatment results in a 99% reduction in signal intensity of each oxonium ion (Figure 2c, bottom panels).

**Figure 2:**
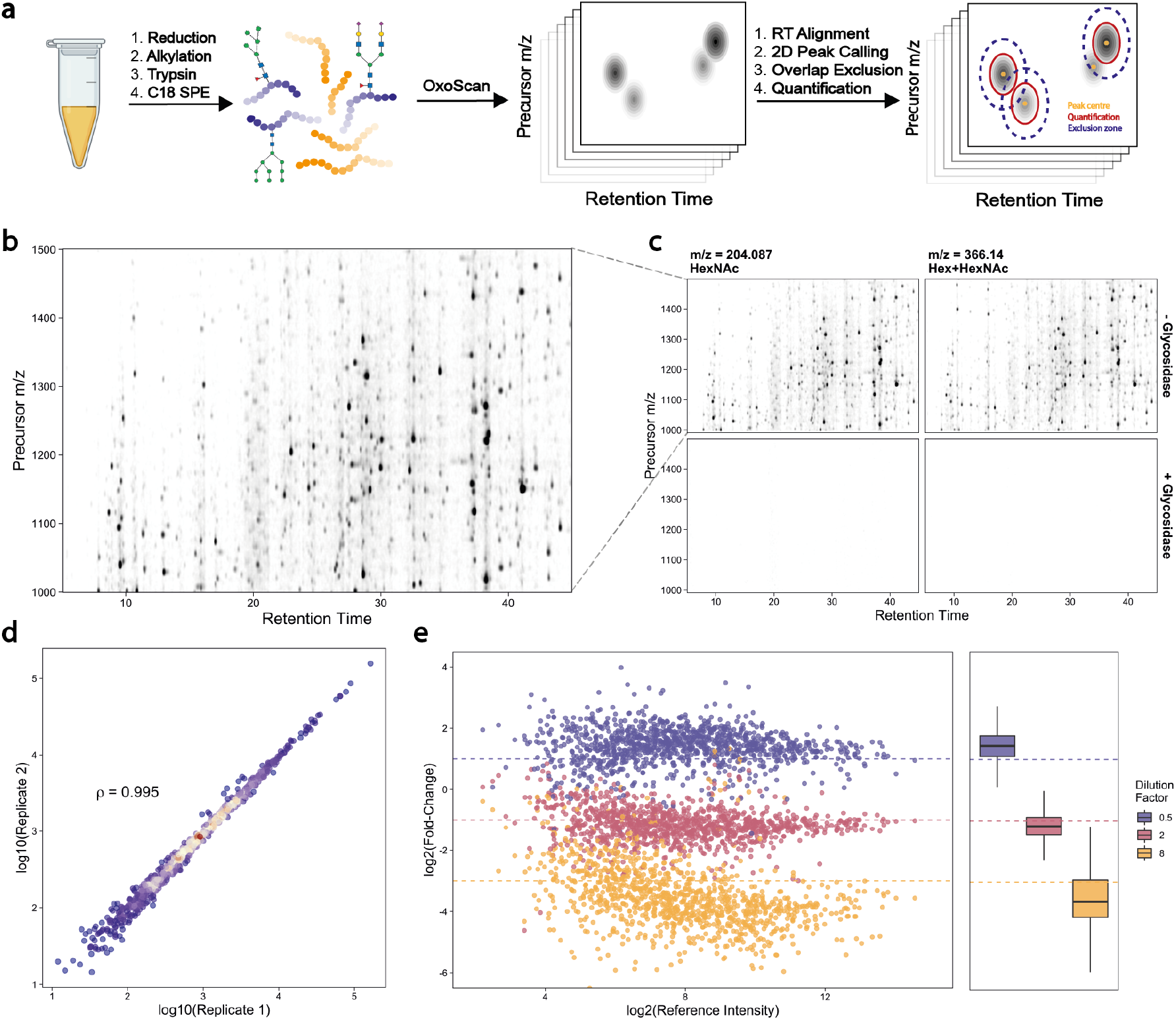
Oxonium ion maps generate a specific and quantitative glycoproteome from the analysis of neat human plasma: **a.** Oxonium ion profiling workflow, starting with the generation of oxonium ion maps from unenriched tryptic digests of serum/plasma (glyco)proteins and computational analysis. The software is freely available at https://github.com/ehwmatt/OxoScan-MS **b.** A glycopeptide map of human plasma tryptic digest, extracted for the m/z = 204.09 (HexNAc) oxonium ion. Each spot represents a glycopeptide in a specific charge state. **c.** Glycopeptide maps for two common oxonium ions present in tryptic digests of human plasma, with and without treatment with a mix of glycosidase enzymes (bottom and top panels, respectively). Peak intensity is proportional to opacity, and all panels are scaled to the maximum peak intensity across the experiment. **d.** Comparison of intensities between two injections of human plasma tryptic digests. Spearman correlation coefficient was calculated based on glycopeptide feature intensities (n = 1006). **e.** Log2(fold-change) plotted for serum glycopeptide features from spiking in 3 different concentrations into an *E. coli* tryptic digest (n = 819). Fold-changes were calculated to a reference dilution factor of 1 and theoretical log2(fold-change) values expected for each dilution factor are plotted as dotted lines. The box-and-whisker plot displays 25th, 50th (median) and 75th percentile in the box. Whiskers display upper/lower limits of data (excluding outliers, not plotted).

To extend this approach for automated and quantitative analysis of oxonium ion profiles, we developed a Python pipeline that utilises a persistent homology-based^60^ algorithm for 2D peak-calling and quantification. For each peak extending into the intensity (*z*) dimension in a glycopeptide map, a ‘persistence’ score is computed, representing the vertical distance between peak maximum and the point where it merges into an adjacent, higher peak. Theoretically, a peak resembling a 2D Gaussian function would have a persistence value equivalent to its height, whereas the persistence value of a peak shoulder would equate to the distance from its apex to the minimum point between the shoulder and the peak (Figure S1d). To facilitate comparison of multiple samples, we implemented retention time alignment using dynamic time-warping.^61^ Upon alignment, peaks are called and ranked by their persistence value. To prevent duplicate calling of a single peak, an exclusion criterion (‘exclusion ellipse’) can be set, within which the centre of another peak with a lower persistence value cannot be called. Quantification is then performed by summing all points in a customisable ‘quantification ellipse’ around each peak maximum. In order to make this analysis approach widely applicable and customisable, Python functions and standalone notebooks with all analysis parameters and requirements are made freely available (https://github.com/ehwmatt/OxoScan-MS).

On neat human plasma tryptic digests, this pipeline identified > 1,100 features, corresponding to glycopeptides in a specific charge state, across over four orders of magnitude in just 19 min of chromatographic separation. The quantities resulting from the 2D peak integration show high reproducibility between replicate injections of a plasma sample (Spearman Correlation Coefficient = 0.994, Figure 2d). We further confirmed quantitative performance by spiking a serum tryptic digest into a background of ^13^C-labelled *E. coli* proteome, maintaining constant total protein content and varying the serum/*E. coli* proteome ratio. Peaks originating from serum features were isolated by removal of any feature identified in a 100% *E. coli* sample. Observed fold-changes in each dilution compared to a reference sample showed agreement with theoretical fold-changes, indicating that differential abundance of glycopeptide features is captured by the OxoScan-MS workflow (Figure 2e).

### The quantitative plasma glycoproteome of severe COVID-19

To test the applicability of OxoScan-MS for cohort studies, we analysed the plasma glycoproteome of a severity-balanced cohort of 30 patients hospitalised due to COVID-19 as well as 15 healthy controls.^18^ Disease severity among patients was assessed according to the WHO Ordinal Scale for Clinical Improvement, ranging from grade 3 (not requiring supplemental oxygen) to grade 7 (requiring invasive mechanical ventilation and additional organ support, Figure 3a). The study protocol and plasma sampling strategies of this cohort was described previously.^18,20^ We utilised microflow chromatography with a 19-minute active gradient and scanned a precursor range optimised for glycopeptides (800–1400 m/z, Figure S2a). Including blanks and quality-control (QC) samples, a total of 164 glycoproteomic samples were measured in ~3 days of instrument time (Figure 3b). Applying our open-source analysis pipeline to the cohort detected 1,102 unique glycopeptide features across all samples, >90% (1,002) of which were precisely quantified across all clinical samples. To assess quantitative reproducibility of the oxonium ion signatures identified, a coefficient of variation (CV) was calculated for each feature within the triplicate measurements of each sample. Repeated analysis of a pooled plasma sample (‘mass spectrometer QC’) and nine replicates of a commercial plasma standard sample (Tebu-Bio) prepared alongside the clinical samples (‘sample preparation QC’) showed reproducibility across the batch measurements, with median CVs of 14% and 20%, respectively. Importantly, the signal as recorded in clinical samples (median CV = 44%) was much higher than this technical variation, indicating that our method detects the biological differences (Figure 3c). The dynamic range of quantified features spans over four orders of magnitude (Figure 3d). 230 glycopeptide features were found to be significantly differentially abundant in response to severe SARS-CoV-2 infection (Figure S2d, log2(FC) > 1, adjusted *P*-value < 0.05, Benjamini–Hochberg multiple testing correction). Consistent with the differential expression analysis, principal component analysis (PCA) and hierarchical clustering show glycoproteomic profiles correctly clustered the majority of healthy/COVID patients (Figure 3e,f), indicating glycopeptide abundance is differentially regulated with increasing COVID-19 disease severity. For three COVID-19 patients, we observed clustering with healthy controls, one of which is explained by very mild disease. It is worth noting, however, that we observed this on both protein- and glycopeptide level.^20,62^

**Figure 3:**
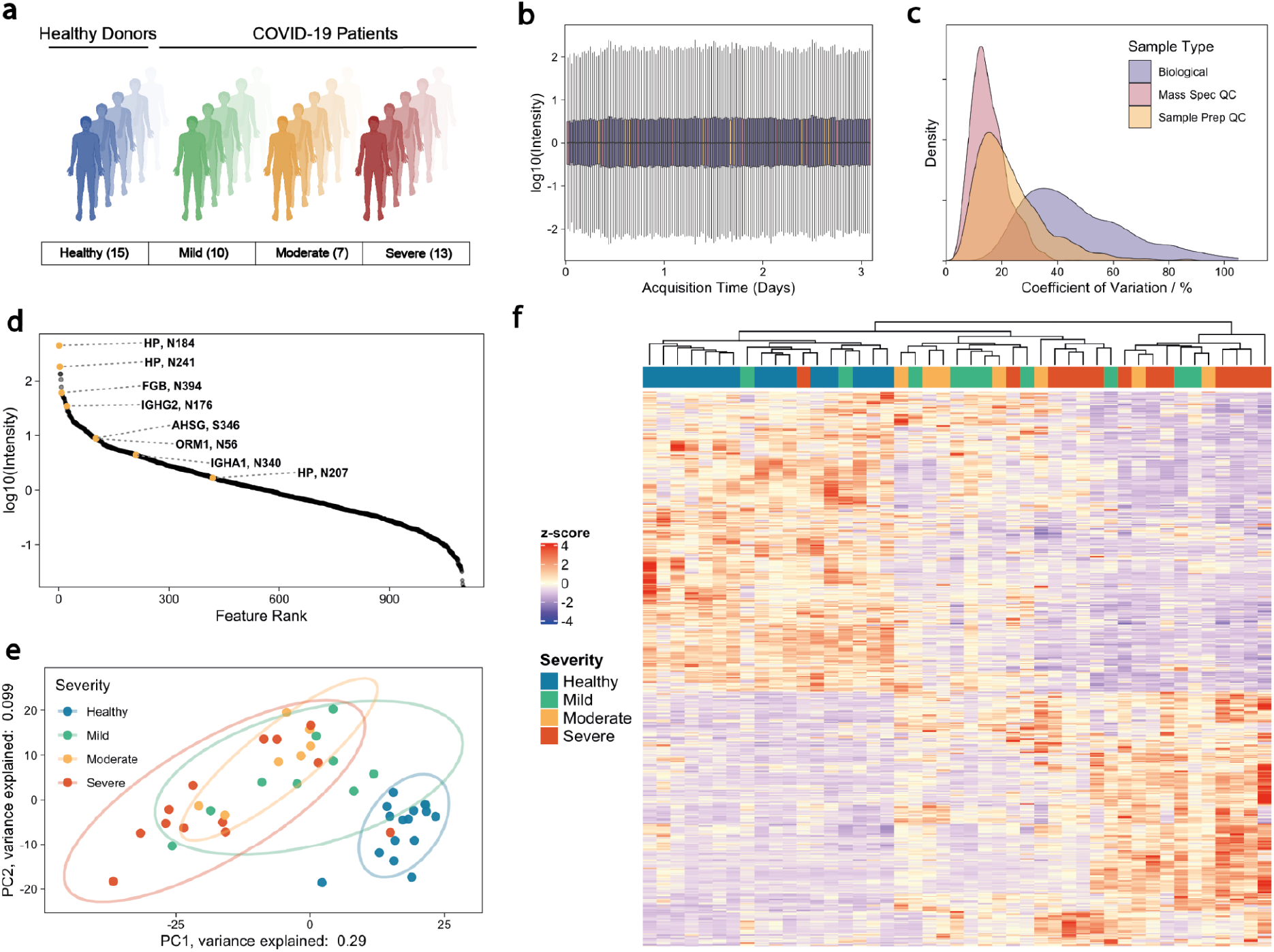
Oxonium ion profiling allows robust and reproducible plasma glycoproteomics in a COVID-19 inpatient cohort: **a.** COVID-19 inpatient cohort, comprising 30 patients hospitalised due to PCR-confirmed SARS-COV2 infection and 15 healthy controls. COVID-19 patients were distributed across different disease severities, ranging from mild (WHO 3), moderate, (WHO 4, 5) and severe (WHO 6, 7) COVID-19. **b.** Total sample intensities across the MS measurement batch following median normalisation; outliers are not plotted. Colour code is identical to panel **c.** Technical and biological variation across cohort measurements, indicated by distributions of coefficient of variation (CV) values for glycopeptide features in repeat injections (Mass Spec QC, *n* = 10), commercial plasma (Tebu) prepared in parallel with samples (Sample Prep QC, *n* = 9), and patient samples (*n* = 3 for each of 45 subjects). **d.** Median intensity of glycopeptides identified in a pooled sample, showing quantification spanning more than four orders of magnitude. **e**. Principal component analysis (PCA) of all robustly detected features (*n* = 1,002) separates healthy and COVID-19 patients in PC1. The proportion of variation accounted for by each axis is shown in axis labels. **f.** Hierarchical clustering of differentially expressed glycopeptide features (calculated using limma R package, adjusted *P*-value < 0.05) between COVID-19 patients and controls.

In order to test glycopeptide assignment after quantification by OxoScan-MS, we selected 26 glycopeptide signals indicative of differential glycan regulation in COVID-19 (Table S5). We then applied a narrow-window and MS1-optimised method to extract accurate precursor masses from pooled cohort samples. Overlay of ‘Q1 profiles’ and MS1 spectra identified high-resolution precursor masses for OxoScan features (Figure 4b). Next, we validated and assigned glycan and peptide information to the differentially-abundant glycopeptides using a complementary fragmentation technology and software, analysing pooled samples by HCD-pd-ETD on an Orbitrap Eclipse (Thermo Fisher Scientific). Following glycopeptide identification with Byonic^63^ (Protein Metrics Inc.) and post-processing filtering for assignment quality (Byonic Score >150, |Log Prob| > 3, as discussed in ^44,64^), glycopeptide identifications were mapped to candidate precursor masses obtained by OxoScan-MS. We then validated the selected glycopeptides based on precursor matching, retention time and comparison of respective DDA- and narrow-window DIA-derived MS/MS spectra (Figure 4c, 4d).

**Figure 4:**
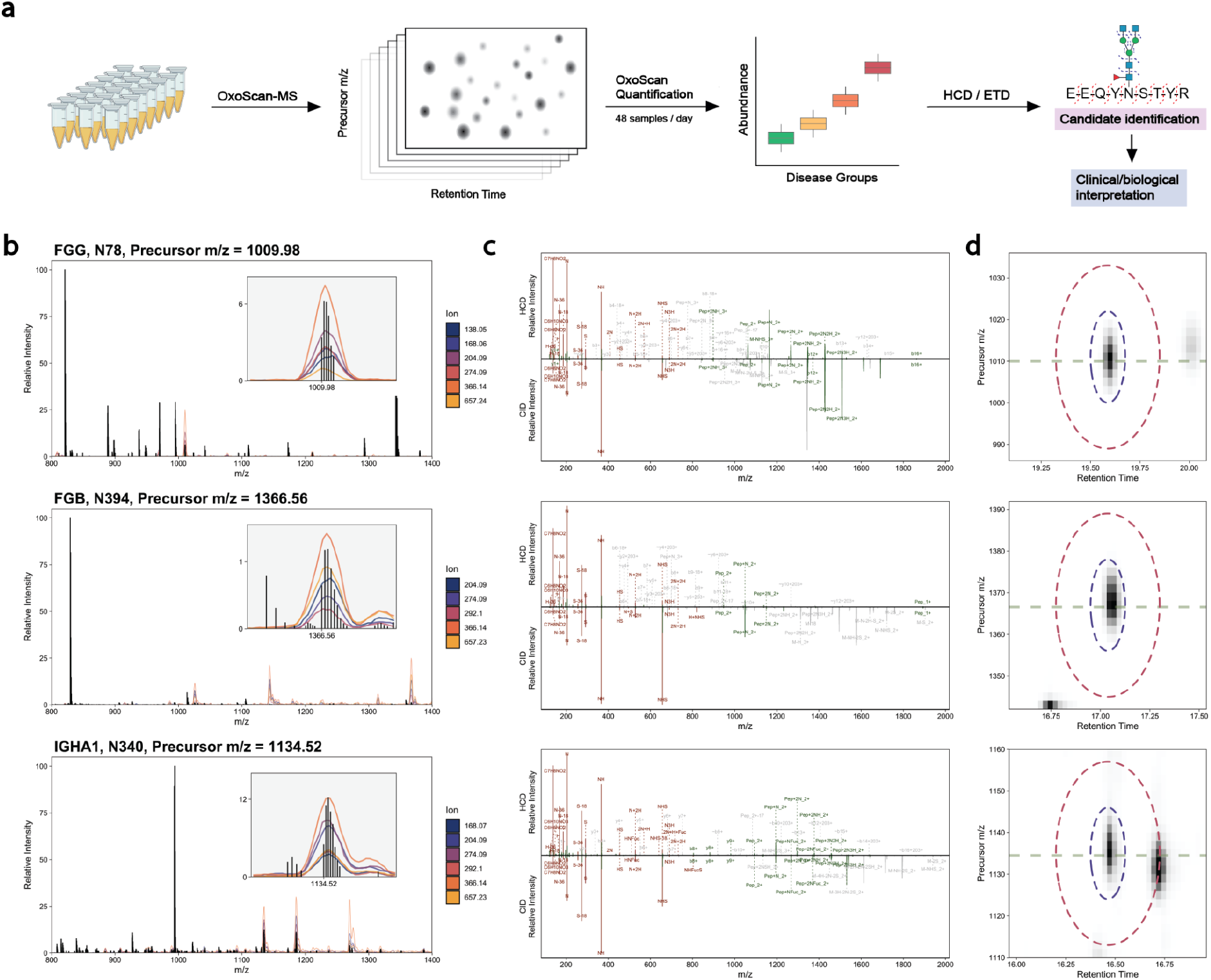
Precursor assignment from the MS1 scanning dimension and subsequent MS/MS matching allows identification of candidate biomarker glycopeptides: **a.** Plasma samples are measured using OxoScan-MS to generate glycopeptide maps and features are identified with complementary fragmentation and database searching. OxoScan then allows quantification of identified features across cohorts with >100s of samples **b.** MS1 spectrum of tryptic plasma digest with Q1 profiles of oxonium ions overlaid. Oxonium ion traces localise glycopeptide precursors even in the presence of co-eluting unmodified peptides of significantly (10-fold) higher abundance. Inset shows zoomed in oxonium ion traces with precursor m/z labelled on x-axis. Q1 profiles were acquired with a 2 m/z scanning window. Vertical panels show fibrinogen gamma chain N-glycopeptide (Asn78), fibrinogen beta chain N-glycopeptide (Asn394) and immunoglobulin A N-glycopeptide (Asn340) respectively. **c.** Comparison of DDA (HCD) and DIA (CID) MS/MS spectra for respective MS1 precursors. Fragment assignments are taken from analysis of DDA data in Byonic (with a tolerance of 5 ppm for DDA and 0.1 Da for DIA). Fragments observed in both DDA and DIA spectra (also matched to within 0.1 Da) are shown in green and oxonium ions are shown in red. All non-matched assignments are shown in grey. Respective panels show the same glycopeptides as in **b**. **d.** Oxonium ion elution profiles in both precursor *m/z* and RT space for respective glycopeptide precursors. Blue and red ellipses represent the quantification and exclusion regions, respectively, and the horizontal line indicates accurate (TOF) precursor mass. Panels show the same glycopeptides as in **b,c**.

This approach identified the 26 selected glycopeptides reflective of differential protein glycosylation in COVID-19, belonging to 11 protein groups (Figure S3, Table S5), These proteins include acute-phase proteins (fibrinogen, haptoglobin, alpha-1-antitrypsin), immunoglobulins (IgG, IgA), and iron-binding proteins (transferrin, hemopexin). A glycopeptide signal could change because of differential glycosylation, a concentration change in the glycosylated protein, or both. To distinguish these cases, we integrated glycopeptide data with protein abundance measurements.^18,20^ On the one hand, we found glycopeptide features that follow the abundance changes of the glycosylated protein. For instance, N- and O-glycopeptides of alpha-2-HS-glycoprotein and N-glycopeptides of hemopexin responded in a similar manner as respective protein abundances (Figure S2e). On the other hand, other signals revealed cases in which the glycan composition or abundance changed differently to the protein abundance, indicating differential glycosylation. For instance, IgA was differentially glycosylated at two separate glycosites, Asn144/131 and Asn340. We observed multiple IgA glycopeptides and note good agreement with previously reported glycopeptide compositions from these sites (% fucosylated, galactosylated, bisecting GlcNAc and sialylated glycans).^57,65^ At Asn 144/131, abundance of smaller, non-sialylated (N2H5, N5H3) glycans increased in accordance with protein abundance changes, whereas monosialylated complex-type biantennary glycans with and without bisecting GlcNAc (N4H5S1, N5H5S1) significantly decreased with increasing COVID-19 severity, indicating potential glycan-specific regulation (Figure 5c). Furthermore, at Asn340, abundance of the disialylated (N5H5S2F1) glycopeptide increased with protein abundance, however the corresponding singly sialylated glycopeptide (N5H5S1F1) showed no significant response in varying disease groups (Figure 5d). Interestingly, IgA levels have been reported to correlate with disease severity in COVID-19 patients and distinct roles identified for each IgA subclass (and their distinctive glycosylation profiles) in inflammation-associated effector function, making a thorough understanding of IgA glycosylation in COVID-19 disease progression highly relevant.^65,66^

We also quantified glycopeptides at two known N-glycosites of alpha-1-antitrypsin (SERPINA1) with biantennary disialylated (N4H5S2) glycans.^67,68^ Although bearing the same glycan modification, we observed an increase of the Asn74 glycopeptide abundance with COVID-19 severity, but a significant decrease of the glycopeptide at Asn271 (Figure 5a). While only a fraction of alpha-1-antitrypsin glycosylation, these results suggest potential regulation of glycan micro-/macro-heterogeneity in COVID-19 patients. Furthermore, aberrant glycosylation of alpha-1-antitrypsin has been observed in the serum of COVID-19 patients by 2-dimensional gel electrophoresis^69^ and in the related alpha-1-antichymotrypsin following septic episodes.^9^

Finally, we detected differential glycosylation of the fibrinogen beta (FGB) and gamma (FGG) chains. Although both protein abundances increase with COVID-19 disease severity, we find the non-sialylated biantennary glycan (N4H5) did not vary significantly between healthy and severe patients at Asn394 of FGB or Asn78 of FGG. Conversely, the mono-sialylated glycan (N4H5S1) was significantly reduced at FGB Asn394, however at FGG Asn78 showed a significant increase (Figure S3), reflected also in the di-sialylated biantennary glycan at this site. These results indicate both site- and glycan-specific regulation of fibrinogen glycosylation in COVID-19 progression. Due to the importance of fibrinogen in COVID-19, these results warrant further investigation to understand the biological cause and consequences of differential fibrinogen glycosylation.

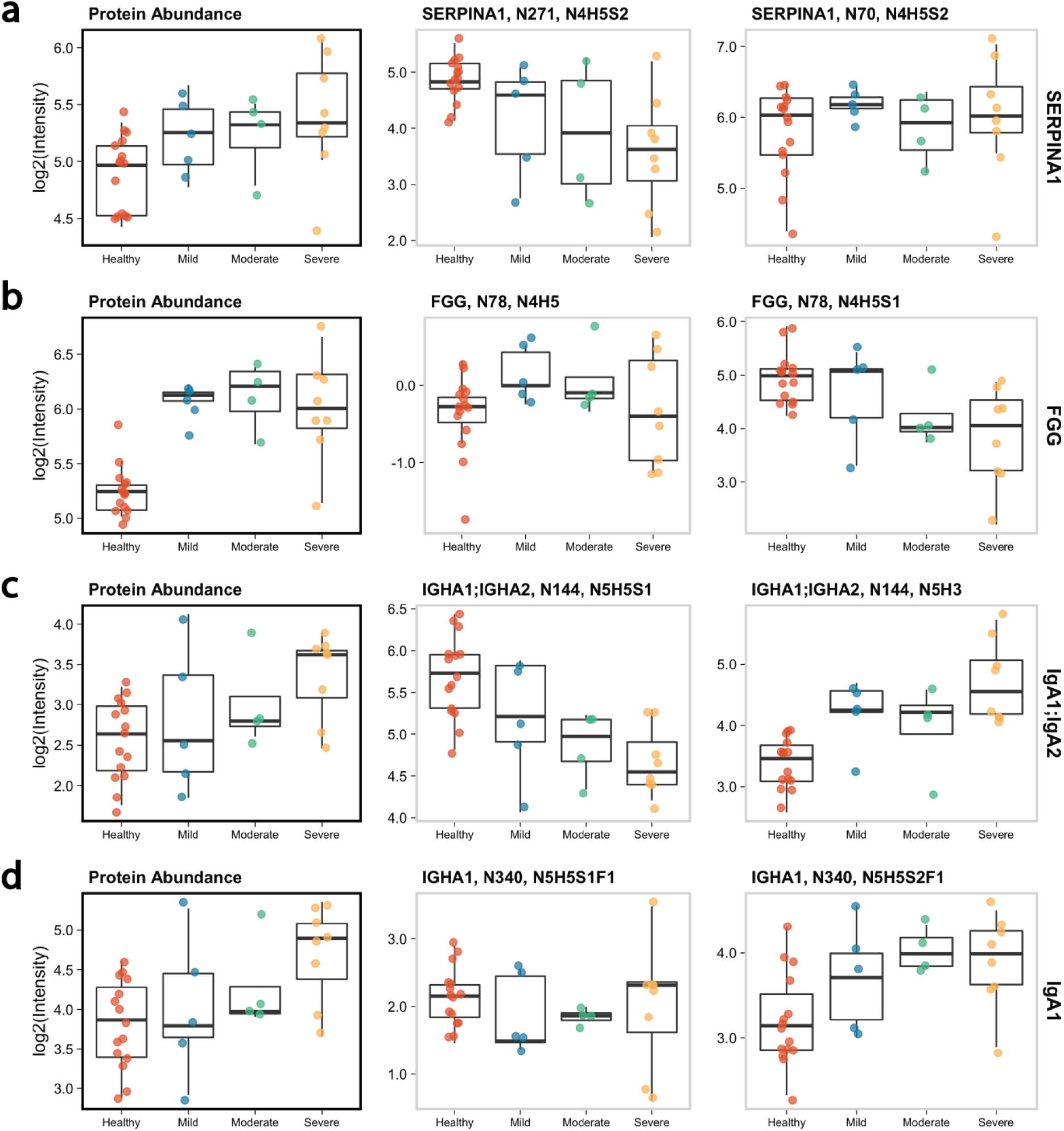
Differential glycosylation of plasma proteins with COVID-19 disease severity identifies potential glycopeptide biomarkers. **a.** Detection of site- and glycan-specific regulation in the plasma glycoproteome of SARS-CoV-2 patients/healthy controls, which cannot be measured by current protein-level analyses. Boxplots show protein (left-most boxplot for each row^20^) and glycopeptide abundances for: **a.** Two different glycosites of alpha-1-antitrypsin (SERPINA1), each bearing a biantennary disialylated (N4H5S2) glycan. **b.** Fibrinogen gamma chain (FGG) glycopeptides contrasting protein abundance changes with increasing disease severity **c.** Immunoglobulin A (IgA, shared glycosite between subclasses 1 and 2) **d.** Immunoglobulin A (IgA, subclass 1 specific glycosite). Relative intensity is scaled to median cohort intensities for both protein and glycopeptide abundances. Boxplots display 25th, 50th (median) and 75th percentile, whiskers display upper/lower limits of data.

## Discussion

Recent studies have attributed substantial potential in the identification of next-generation glyco-biomarkers and predictive signatures,^70–72^ but due to the complexity of protein glycosylation, large scale analysis of plasma and serum glycosylation remains a major challenge. Here we present OxoScan-MS, and demonstrate robust and reproducible quantification of over 1,000 glycopeptide features in neat plasma, with a total run-time per sample of less than 30 minutes and no requirement for glycopeptide enrichment. OxoScan-MS operates by scanning for and quantifying diagnostic oxonium ions, followed by targeted feature identification. OxoScan-MS is hence not a replacement for current glycoproteomic techniques; it is rather complementing them with a fast, quantitative and highly cost-effective method to screen large cohorts. In contrast to DDA-based glycopeptide approaches where the co-elution of unmodified peptides reduces the time spent analysing glycopeptides specifically, OxoScan-MS samples glycopeptides independently of co-eluting unmodified peptides and is therefore compatible with samples prepared for protein-level analyses, combining the advantages of a precursor ion scan with SWATH-MS to provide a digital snapshot of the glycoproteome. To our knowledge, OxoScan-MS is thus the first technology which allows glycoproteomic profiling of hundreds of samples prepared for conventional MS-based proteomics.

We applied OxoScan-MS to study the plasma glycoproteome in response to SARS-CoV-2 infection, measuring a severity-balanced clinical inpatient cohort in triplicate (164 samples in total) in just 3 days of instrument time. From the glycopeptide features measured, 230 were differentially concentrated between healthy and severely affected patients. We then selected 26 of these features and determined their identity using conventional glycoproteomic approaches. We found altered glycopeptide abundances among proteins important in COVID-19, including haptoglobin, hemopexin, fibrinogen, immunoglobulins A and G. Furthermore, by integrating protein abundance data we identify glycan-specific regulation dependent on disease severity, most notably for alpha-1-antitrypsin, IgA, and fibrinogen. These three proteins are involved in inflammation, immune response, and coagulation, three key systems in the pathogenesis of COVID-19.^73^ Reassuringly, alpha-1-antitrypsin, IgA, and fibrinogen are indicators of COVID-19 disease severity^74,75^ and hence, our results associated their differential glycosylation to severe COVID-19.

We anticipate that large-scale clinical glycoproteomic profiling, supported by increasingly high-throughput and quantitative glycoproteomics technologies, can aid in the discovery of glycoform-specific biomarkers, relevant for understanding disease mechanisms as well as for diagnosis, prognosis and prediction. No enrichment steps were used in the presented study, enabling a robust workflow for clinical applications where reproducibility is of utmost importance. However, we would like to emphasise that the dynamic range and depth might be further increased by removing highly abundant proteins and unmodified peptides with a variety of enrichment strategies or increased acquisition times per sample. This is a common trade-off in (glyco)proteomics experiments, however for our purposes we focused on increasing the practical throughput and costs of glycoproteomics experiments. We also note that although we identify both N- and O-glycopeptides, further developments could improve the detection of O-glycosylated peptides. Beyond plasma biomarker discovery, we anticipate OxoScan-MS could have a number of immediate applications, for example to high-throughput glycoprofiling of biologics and the workhorse cell lines used to produce them. As the reported data processing and analysis methods are ion-agnostic, our workflow can also be applied wherever characteristic fragment ions are produced, such as for other endogenous post-translational modifications^76,77^ or rationally designed chemical probes.^78^

## Materials and methods

### Materials

LC-MS grade reagents were purchased as follows: water (Fisher Scientific, PN: 10505904), acetonitrile (ACN, Fisher Scientific, PN: 10001334), methanol (MeOH, Fisher Scientific, PN: 10767665), formic acid (FA, Pierce, PN: 85178), trifluoroacetic acid (TFA, Sigma-Aldrich, 85183), DL-dithiothreitol (DTT, Sigma-Aldrich, PN: 43815), iodoacetamide (IAA, Sigma-Aldrich, PN: I1149), urea (Sigma-Aldrich, PN: 1084870500), ammonium bicarbonate (ABC, Fisher Scientific, PN: 15645440). Trypsin was purchased from Promega (PN: V5117). Solid-phase extraction plates were purchased from NEST Inc. (BioPureSPN Macro 96-Well, 100mg PROTO 300 C18, PN: HNS S18V-L).

### IgG Isolation from Human Serum

IgG was purified from human serum samples as described previously.^55^ In brief, IgG was isolated from 5 μl of serum using 30 μl of Protein A Sepharose (GE Healthcare, Eindhoven, The Netherlands). Sample mixtures were incubated under agitation at 650 rpm for 1 hour at room temperature. Protein A Sepharose beads were washed with 5×200 μl 1×PBS and 3×200 μl MilliQ water. IgG were eluted with 3×100 μl of 100 mM formic acid. Eluates were dried in vacuum centrifuge then redissolved in 50 μl of 50 mM ammonium bicarbonate and shaken for 5 min. Sequencing grade trypsin (Promega, Madison, WI) was added to a final concentration of 0.2 μg/μl and samples were incubated overnight at 37 °C. On the following day, IgG glycopeptides were isolated from peptides using self-made micro-spin cotton-HILIC columns. They were conditioned by washing with 3×50 μl MilliQ water and 3×50 μl 80% ACN. Afterwards, dried IgG samples were resuspended in 50 μl of 80% ACN and loaded on the self-made microcolumns. They were washed with 3×50 μl 80% ACN containing 0.1% TFA and then with 3×50 μl 80% ACN. The retained IgG glycopeptides were eluted with 6×50 μl MilliQ water, dried out in a vacuum centrifuge and stored at −20 °C until measurement.

### Standard preparation of IgG and serum samples

20μg purified IgG or 5μl of raw plasma/serum were prepared as previously described.^20^ In brief, IgG/plasma was denatured and reduced by addition of 55μl 8M Urea, 5.5mM DTT, 100mM ABC and incubation for 1h at 30°C. All subsequent steps were carried out using a Beckman Coulter Biomek NXP 96-well liquid handling robot. 5μl of 100mM IAA was added and incubated in the dark for 30 minutes. Reduced/alkylated proteins were then diluted with 340μl 100mM ammonium bicarbonate (to bring [Urea] < 2M) and digested with trypsin (1:50 w/w) for 17h at 37°C. Digestion was stopped by acidification with 25μl 10% formic acid and peptides were cleaned up by solid-phase extraction (NEST C18 MacroSPIN SPE plates, as described previously18). In brief, each well was treated/centrifuged sequentially in the following steps: 200μl MeOH, 1min @ 50g, 2 x 200μl 50% ACN, 1min @ 150g, 2 x 200μl 0.1% FA, 1min @ 150g, 200μl sample, 1min @ 150g, 2 x 200μl 0.1% FA, 1min @ 200g, 1min @ 200g, 3 x 110μl 50% ACN, 1min @ 200g. Elution (50% ACN) fractions were eluted into the same respective wells and dried in an Eppendorf Speedvac (45°C, ~7h). Dried, desalted peptides were resuspended in 0.1% FA (0.5 - 2μg/μl depending on sample) and stored at −80°C until measurement.

### Glycosidase Treatment

Deglycosylation was performed with the Protein Deglycosylation Mix II (New England Biosciences, PN: P6044S). For glycosidase treatment, plasma samples were prepared as described above with the following modifications: Following dilution of reduced/alkylated plasma with 340μl 100mM ABC, 45μl 10X Protein Deglycosylation Buffer I was added. Next, 5μl of either Protein Deglycosylation Mix II (New England Biosciences, PN: P6044S) or 100mM ABC (for deglycosylation and control respectively) were added and incubated at RT for 30 minutes and at 37°C for a further 16h. Following deglycosylation, tryptic digest and SPE was performed as described above. Dried samples were redissolved in 50μl 0.1% FA and injected as is. Samples were measured with a 45-minute water-to-acetonitrile gradient with a 10 m/z Scanning SWATH window (see Table S4).

### Heavy-labelled *E. coli* growth and sample preparation

*E. coli* MG1665 was woken on LB agar and grown in M9 minimal media supplemented with ^13^C-glucose (11.28g/L M9 salts, 2mM MgSO_4_, 0.1mM CaCl_2_), 1% ^13^C-glucose). Cells were harvested at mid-log phase, washed with water and lysed in 200μl 7M Urea, 100mM ABC with acid-washed glass beads (425-600 μm). Samples were then prepared as described previously18. Briefly, cells were lysed with mechanical bead beating (1600 MiniG, Spex Sample Prep) for 5 min at 1500rpm, reduced with 20μl 55mM DTT for 60 min at 30°C and subsequently alkylated with 20μl 120mM IAA at room temperature in the dark for 30 minutes. Lysates were then diluted with 1ml 100mM ABC, centrifuged at 3220g for 5 minutes and the supernatant taken for tryptic digest (9μl of 0.1μg/μl solution) for 17h at 37°C. Acidification and SPE clean-up was performed as described for plasma, with the following modifications: 3% ACN, 0.1% FA was used instead of 0.1% FA and elution volumes were 120μl, 120μl and 130μl. Eluted peptides were dried and redissolved as described for plasma.

### Spike-in sample preparation

Commercial serum tryptic digests (prepared as described above) and heavy-labelled *E. coli* tryptic digests were resuspended in 0.1% FA and peptide concentration measured on a Lunatic spectrophotometer. The digests were subsequently mixed in set ratios by protein amount (serum*:E. coli;* 5:95, 20:80, 40:60, 80:20), normalised to the same sample volume and 2μg injected for each sample.

### COVID-19 Cohort Analysis

Patient samples were prepared as described in the general workflow and processed without further enrichment/depletion. The 45 biological samples were randomised into 96-well plate format and prepared in whole-process triplicate alongside aliquots of commercial plasma citrate. In order to minimise the effect of instrument drift, samples were block randomised by replicate for sample acquisition. A pooled plasma sample was generated by mixing a small aliquot of tryptic peptides from each clinical sample (MS.QC, *n* = 10) and measured every 16 samples throughout the batch to monitor instrument performance. Commercial plasma was added to 96-well plates and prepared in parallel with the clinical samples as whole-process QCs (SP.QC, *n* = 9). Blanks and mass calibration samples (“Pepcal”) were also included every 16 injections across the cohort.

### Data-independent acquisition (OxoScan-MS)

All Scanning SWATH/DIA analysis was performed on a Waters NanoAcquity coupled to a Sciex TripleTOF 6600. Peptides were separated on a reverse-phase C18 Waters HSS T3 column (1.8μm, 300μm x 150mm, 35°C column temperature) at 5μl/min (loading flow/buffers). Peptides were separated with gradients of buffer A (1% acetonitrile, 0.1% formic acid) and buffer B (acetonitrile, 0.1% formic acid). The Cohort method ramped with a non-linear gradient from 3-40% B over 19 minutes (Table S3), while glycosidase treatment and GPF chromatographic gradients ramped linearly from 3-40% over 45 and 90 minutes respectively. For IgG analysis, a linear gradient ramped from 3-18% Buffer B over 90 minutes. Upon reaching 40% in the respective gradients, washing and re-equilibration steps were as follows: 40-80% B over 1 minute, 80% B for 0.5 minutes, 80-3% B over 1 minute, re-equilibration at 3% B for 6 minutes until next injection. Source conditions were as follows: Source Gas 1: 15 psi, Source Gas 2: 20 psi, Curtain Gas: 25 psi, Temperature: 0°C, IonSpray Floating Voltage: 5500 V, Declustering Potential: 80 V. Rolling collision energies were calculated from the following equation: CE = 0.034 * m/z + 2, where m/z is the centre of the scanning quadrupole bin. Precursor range, window width and cycle times were tailored depending on chromatographic gradient, desired Q1 resolution and sensitivity (Table S4).

### Data-dependent acquisition

Samples were pooled from all healthy and severe patients and analysed on an Orbitrap Eclipse coupled to an Ultimate 3000 RSLCnano (both Thermo Scientific). Sample (1ul, approx. 1μg/μl in 0.1% FA) was loaded onto a trap column (Acclaim PepMap-100 75 μm × 2 cm NanoViper) with loading buffer (2% ACN, 0.05% TFA) at 7 μL/min for 6 min (40°C). Peptides were separated on an analytical column (PepMap RSLC C18 75 μm × 50 cm, 2 μm particle size, 100 Å pore size, reversed-phase EASY-Spray, Thermo Scientific) from 2-40% Buffer B over 87 minutes at 275 nL/min and with the following parameters: column temperature: 40 °C, spray voltage: 2400 V. Gradient elution buffers were: A, 0.1% FA, 5% DMSO and B: 0.1% FA, 5% DMSO, 75% ACN. For MS scans acquired in the Orbitrap, scan resolution was set to 120,000 at FWHM 200 m/z. The precursor range was 400–2000 *m/z* with the following parameters: RF Lens 30%, AGC target 100%, maximum injection time 50 ms, spectra acquired in profile. Monoisotopic peak determination was set to the peptide mode. Dynamic exclusion was enabled to exclude for 10s after *n* = 3 times within 10s with mass tolerance of +/- 10ppm. Precursors (*z* = 2-6) were selected for DDA MS/MS with a quadrupole isolation window of width 2 m/z and a fixed cycle time of 3s. HCD MS/MS scans were acquired in the Orbitrap at a resolution of 30,000 and a normalised collision energy of 28% with the following parameters: First mass m/z 100, AGC target 100%, custom maximum injection time 54 ms, scan data acquired in centroid mode. An HCD-pd-ETD instrument method, whereby ETD fragmentation was only performed if three of the following list of mass trigger ions were present in the HCD MS/MS spectra (±20ppm) and above the relative intensity threshold of 5% (126.055, 138.0549, 144.0655, 168.0654, 186.076, 204.0855, 366.1395, 292.1027, 274.0921, 657.2349 *m*/*z*). Precursor priority was given by highest charge state and ETD activation used calibrated charge-dependent ETD parameters, the single scan per cycle was detected in the ion trap with the following parameters: isolation window of 3 *m/z,* rapid scan rate, first mass m/z 100, AGC target 100%, custom maximum injection time 54 ms, scan data acquired in centroid mode.

### DIA Data processing

Raw Scanning SWATH data files (.raw) were processed to Sciex .wiff format using the Scanning SWATH Raw Processor (AB Sciex) using the default settings except for the following: Q1 binning = 4. Wiff files were then converted to .dia files in DIA-NN and XICs were extracted (as .txt files) across the entire precursor range using the --extract [oxonium ion masses] function. The output text files were directly imported into OxoScan scripts (as a Jupyter Notebook). For the COVID-19 cohort method, the following settings were used: Maximum number of features called = 5000, m/z bin width = 2 (*m/z*), RT bin width = 0.025 min, m/z quantification radius = 5 (bins), RT quantification radius = 3 (bins), m/z exclusion radius = 2 * m/z quantification radius and RT exclusion radius = 3 * RT quantification radius. Samples were normalised and scaled prior to retention time alignment to prevent distortions due to sample intensity, but output quantities were not normalised.

### Data Analysis

All processed data (OxoScan/Byonic output, exported MS data) was analysed using custom R scripts. General data manipulation was carried out with tidyverse packages^79^ and visualisation with ggplot2.^80^ Differential expression analysis was performed with the limma R package^81^ and Kendall-Tau trend test (as part of the EnvStats package^82^). Heatmaps were plotted with the ComplexHeatmap R package.^83^ Principal component analysis was carried out with the prcomp R package. PeakView (AB Sciex) was used for accessing raw MS data for precursor mass assignment, manual inspection and exporting of spectra/XICs.

All analysis scripts and figure generation can be reproduced at https://github.com/ehwmatt/OxoScan-MS. In brief, for each patient, a mean sample intensity and CV was calculated for each feature from three technical replicates and used for further analysis/statistical testing. 5 samples were removed from the analysis due to low signal intensity and all samples were median normalised. In order to prevent misidentification of non-glycosylated precursors by interfering signals in the oxonium ion regions, features for which a single oxonium ion comprised >85% of the total oxonium ion signal were removed. Furthermore, specific ion signals were removed if the percentage contribution for a given feature showed significant variability (indicating interference/poor quantitation). Finally, features were kept for quantification only if >3 oxonium ions were quantified across all samples in the clinical cohort.

### DDA Data Processing

Data-dependent glycoproteomics experiments were analysed in Byonic (Protein Metrics Inc., v4.1.5).^63^ Thermo raw files were searched against the Uniprot Human FASTA (3AUP000005640-canonical, downloaded 26^th^ May 2018) and a built in library of 57 human plasma glycans, 132 human N-glycans and 9 human O-glycans, all set as ‘rare1’. Carbamidomethylation (+57.0214) was set as a fixed modification and oxidation (+15.9949) as ‘common1’. Tryptic digest was selected (RK, ‘C-terminal cutter’, fully-specific, max. 1 missed cleavage). The following search parameters were applied, precursor tolerance: 5 ppm, fragment tolerance: (HCD) 5 ppm, fragment tolerance (ETD): 0.6 Da, Protein FDR, 1%. Identified glycopeptide information (‘Spectra’ tab of each Byonic output file) was imported into R and PSMs were further filtered with the following thresholds: Presence of glycan in ‘Glycans NHFAGNa’ column, Byonic score > 150, |Log Prob| > 3. The resulting identification table was taken forward for matching to identified DIA glycopeptide features with custom R scripts and manual validation, as described below.

### DIA High Resolution MS1 assignment

Glycopeptide precursors were identified in pooled plasma samples using two MS methods (with the same chromatographic gradient and precursor range as the cohort):

1. Q1 method - 2 m/z Scanning SWATH window and total cycle time of 3.6 seconds
2. MS1 method - MS1 scans only with 500ms accumulation time

Precursor masses were identified by extracting oxonium ion chromatograms and Q1 profiles over the RT/binned precursor m/z for specific features (either from a specific ‘peak_num’ in Table S5 or specific glycopeptide identified in DDA experiments) in the Q1 method. For each feature, the reported MS/MS spectra were exported directly for DDA/DIA comparison and fragment assignment. The respective accurate precursor m/z was then extracted in the MS1 method with a tolerance of 0.1 Da and retention times compared. The MS1 spectra were exported directly from PeakView (AB Sciex). High resolution precursor m/z values were used to calculate precursor mass and matched to Byonic-reported glycopeptide precursors with a tolerance of 0.5 Da.

### MS/MS Matching and Glycopeptide Validation

To compare DDA/DIA MS/MS spectra, both HCD spectra and fragment ion assignments from each identified glycopeptide were exported from Byonic as text files. Extracted Scanning SWATH MS and MS/MS spectra (as described above) were exported as text files. Matching fragments were compared between DDA/DIA spectra with a custom R script. For MS/MS matching between DDA/DIA experiments, a list of theoretical and observed fragment ions was exported directly from Byonic for each feature. DDA spectra were matched first to the Byonic fragment list with a tolerance of 0.1 Da and subsequently with the DIA MS/MS spectra with a tolerance of 0.1 Da. In the case of multiple matches, only the match with the lowest mass error was taken.

## Acknowledgements

We thank the organisers and all collaborators at the 2020 Crick Data Challenge, without whom we would not have conceived or developed the OxoScan analysis approach. Figures 2a and 3a were created with BioRender.com. This work was supported by the Francis Crick Institute, which receives its core funding from Cancer Research UK (FC001134), the UK Medical Research Council (FC001134) and the Wellcome Trust (FC001134). Part of this research was funded by the European Research Council (ERC) under grant agreement ERC-SyG-2020 951475, the Wellcome Trust (IA 200829/Z/16/Z). The and by the Ministry of Education and Research (BMBF), as part of the National Research Node ‘Mass spectrometry in Systems Medicine (MSCoresys), under grant agreement 031L0220.

## Author contributions

MEHW, CBM and MR designed the study. MEHW and LK prepared samples for glycoproteomic analysis. MEHW, CBM and HRF carried out mass spectrometry experiments. DMJ, JDF, SKA, MEHW and CBM developed the OxoScan Python analysis package. MEHW, CBM, VD, DMJ analysed the data. PTL and FK collected COVID-19 clinical samples. MEHW, CBM and MR wrote the manuscript, with input from all co-authors.

## Data/code availability

Raw MS data, extracted oxonium ion .txt files from DIA-NN and OxoScan-MS processed outputs have been submitted to MassIVE and will be available on ProteomeXchange upon publication (Accession number: PXD034172). All code (OxoScan Python functions/Jupyter notebooks and R scripts for analysis/reproducing all figures) and OxoScan-MS processed data for IgG, spike-in experiment and the COVID-19 cohort are freely available at https://github.com/ehwmatt/OxoScan-MS.

