## Supplementary Information for "OxoScan-MS: Oxonium ion scanning mass spectrometry facilitates plasma glycoproteomics in large scale"

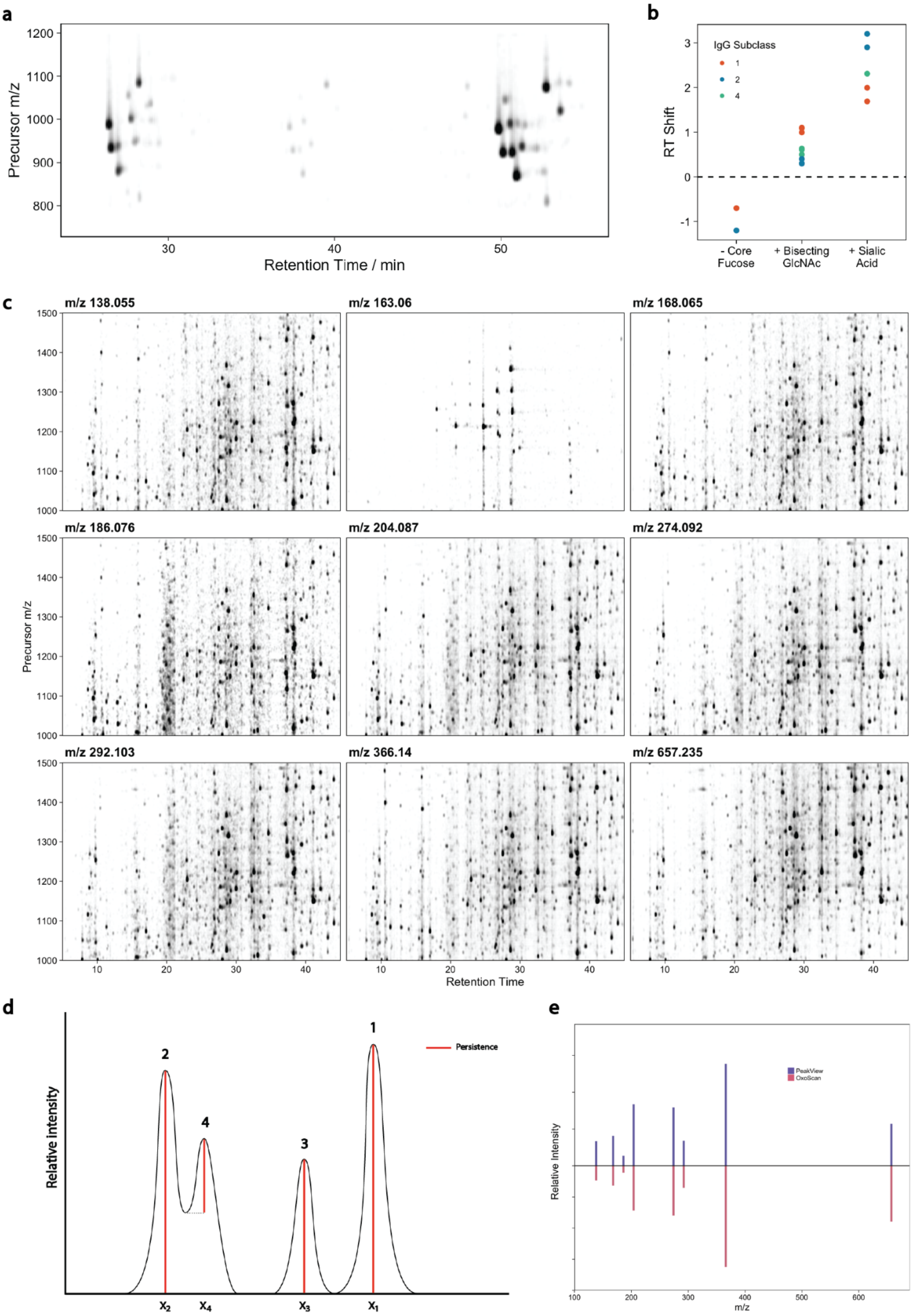


**Figure S1: Qualitative and quantitative glycoproteomic analysis by OxoScan-MS. a.** Oxonium ion map of purified IgG from human serum[1], showing different total abundances of IgG1, 4 and 2 subclasses, from left to right. Oxonium ion signals were extracted in DIA-NN[2], summed and plotted with opacity proportional to intensity. **b.** Retention time shifts in reverse-phase (C18) chromatography of identified IgG glycopeptides upon change of glycan composition, when compared to respective G<u>X</u>F (reference) glycopeptides. **c.** Oxonium ion maps of a human tryptic digest for 9 oxonium ions, extracted in DIA-NN (with a 20ppm mass tolerance) and point opacity plotted proportional to intensity (scaled separately by ion). **d.** Schematic showing the order of priority for peak calling (in 1-dimension) by the persistent homology algorithm. Peak numbering shows rank of persistence values and red lines represent the computed persistence value for each peak. Importantly, peaks are ranked by persistence as opposed to maximum height. **e.** Back-to-back MS/MS spectra of an IgG glycopeptide showing intensities of 8 oxonium ions when exported directly from the MS/MS spectrum (blue, top panel) in PeakView (AB Sciex) compared to output values from OxoScan quantification (red, bottom panel).

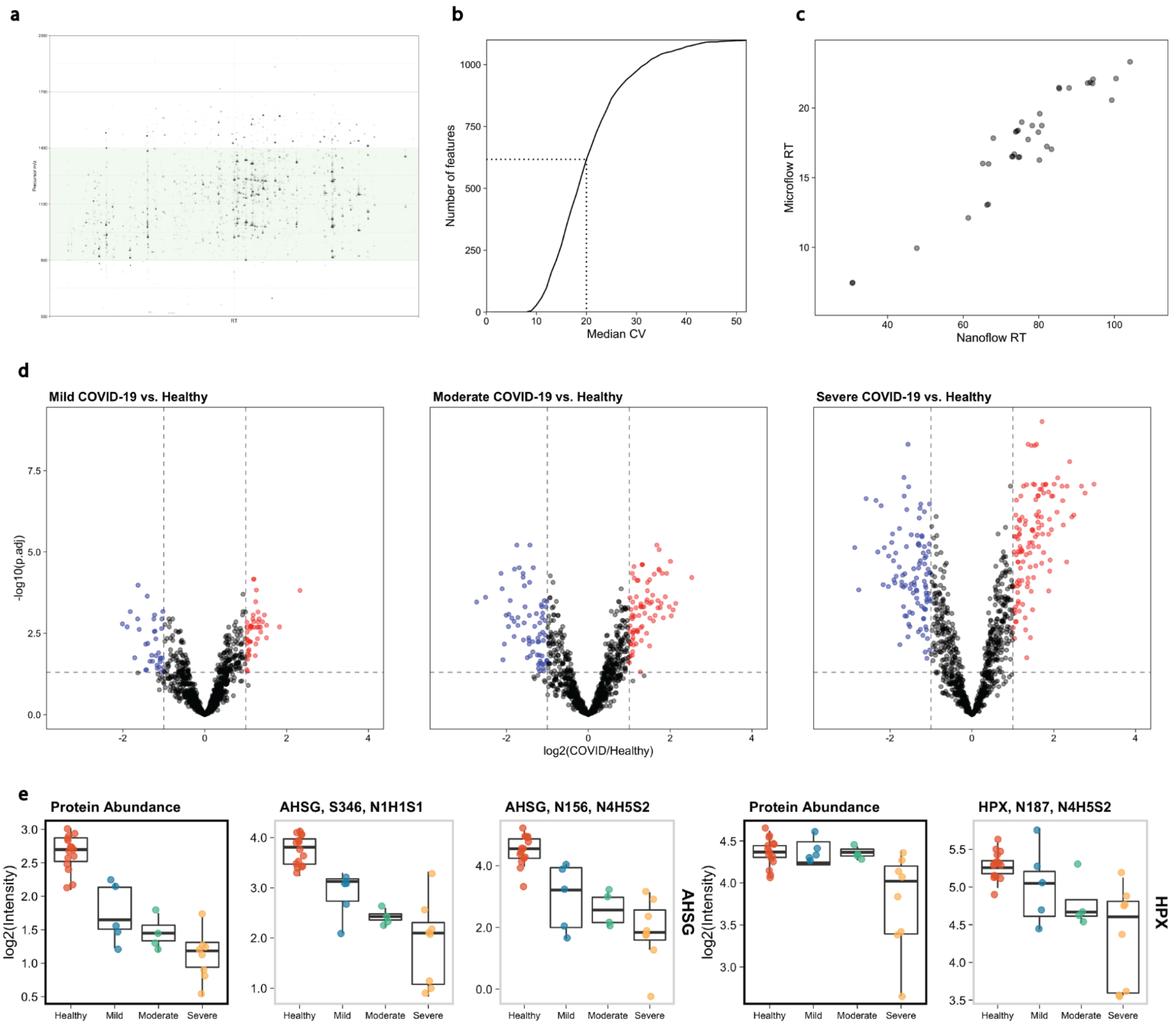


**Figure S2: Profiling the glycoproteomic changes in SARS-CoV-2 infection by OxoScan-MS. a.** Gas-phase fractionation of a single commercial plasma tryptic digest over the precursor range m/z 500-2000 (in 3 separate runs, shown aggregated here) shows the optimum range for detection of glycopeptides by OxoScan-MS. **b.** Median CV (%) values for each feature quantified in clinical samples. CVs were calculated for each feature in triplicate measurements of each patient/donor sample, the median taken for each feature, ranked and plotted against feature number. Dotted line shows the CV = 20% threshold. **c.** Comparison of retention times for glycopeptides identified in both DDA (nano-flow, x axis) and DIA (micro-flow, y-axis) shows good agreement across different chromatographic platforms. **d.** Volcano plots comparing $\log_2$(fold-change) for all glycopeptide features between each grouped disease severity (mild, moderate, severe) against healthy controls. $\text{Log}_2$(fold-change) and p-values were calculated using the limma R package.[3] Multiple testing correction was performed by the Benjamini-Hochberg method.[4] Coloured points represent those with $|\log_2(\text{fold-change})| > 1$ and $P < 0.05$) for up- and down-regulated features (red and blue respectively). **e.** Glycopeptide abundance changes matching parent protein abundance changes with increasing COVID-19 disease severity for alpha-HS-glycoprotein and hemopexin.

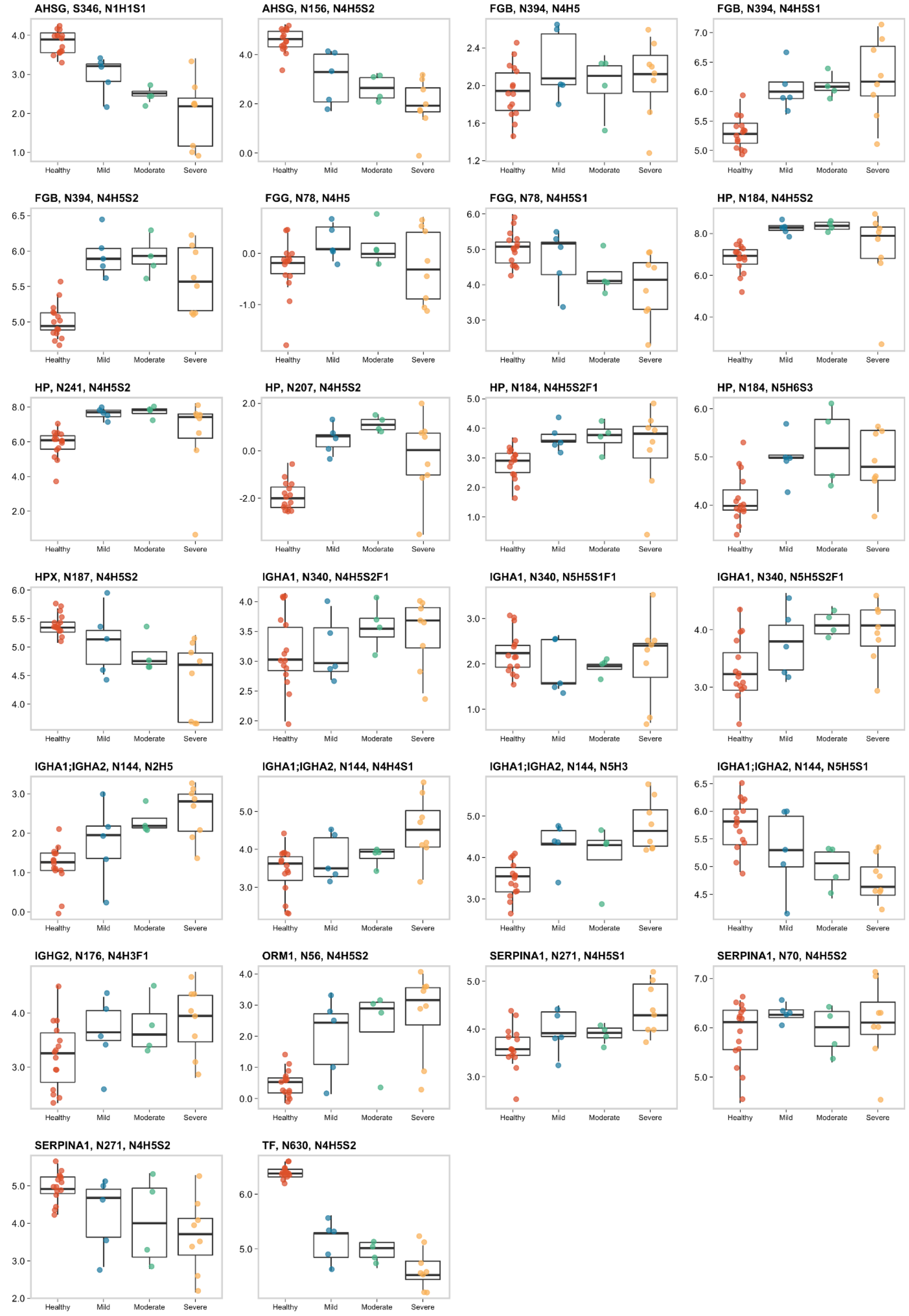


**Figure S3:** Abundances of glycopeptides identified in the COVID-19 cohort, grouped by disease severity. Values are $log_2$-transformed, box-and-whisker plot displays 25th, 50th (median) and 75th percentile in the box. Whiskers display upper/lower limits of data. Plot labels show gene, glycosylation site and glycan composition.

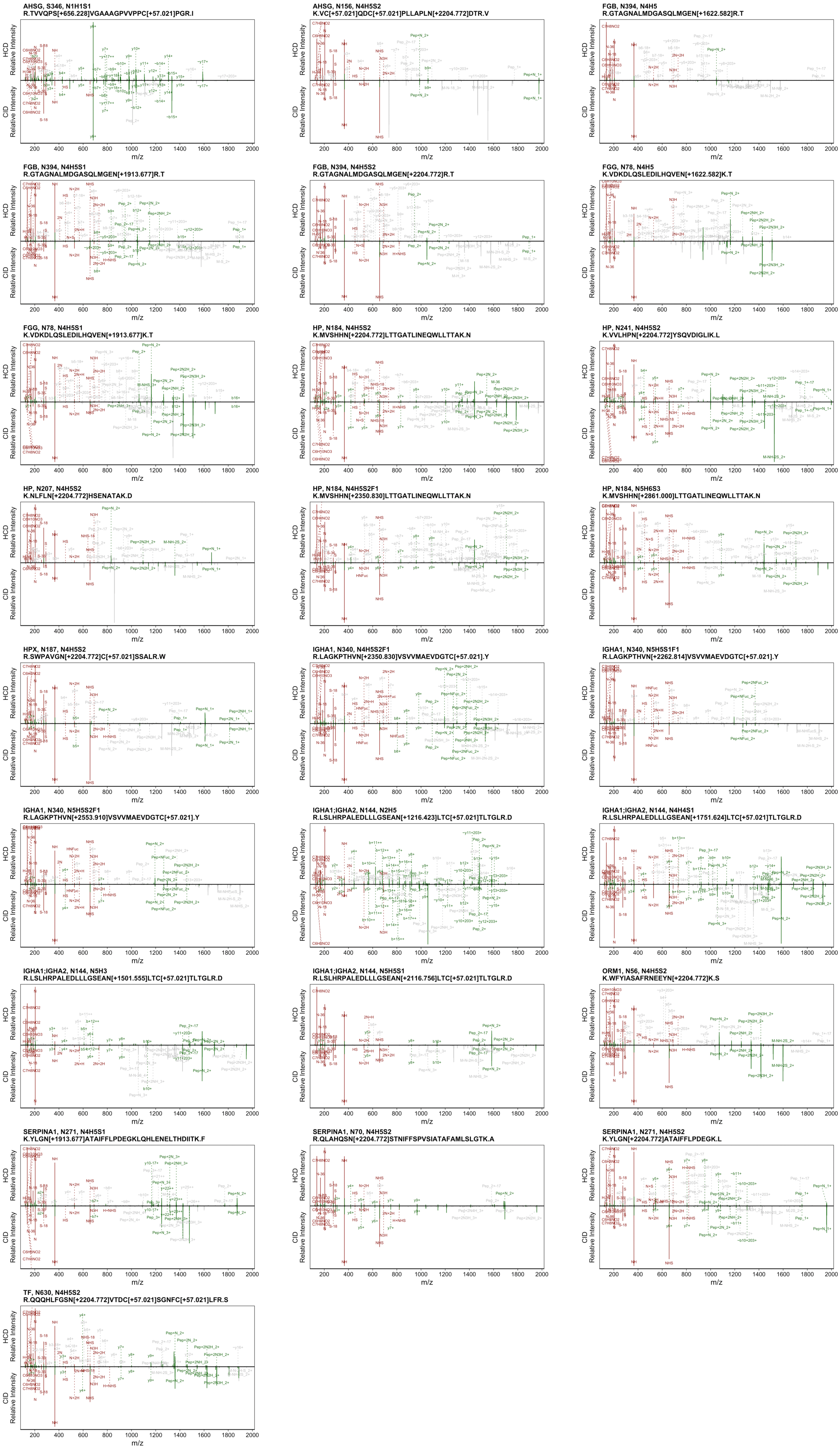
AHSG, S346, N1H1S1
R.TVVQPS[+656.228]VGAAAGPVVPPC[+57.021]PGR.I
AHSG, N156, N4H5S2
K.VC[+57.021]QDC[+57.021]PLLAPLN[+2204.772]DTR.V
FGB, N394, N4H5
R.GTAGNALMDGASQLMGEN[+1622.582]R.T
FGB, N394, N4H5S1
R.GTAGNALMDGASQLMGEN[+1913.677]R.T
FGB, N394, N4H5S2
R.GTAGNALMDGASQLMGEN[+2204.772]R.T
FGG, N78, N4H5
K.VDKDLQSLEDILHQVEN[+1622.582]K.T
FGG, N78, N4H5S1
K.VDKDLQSLEDILHQVEN[+1913.677]K.T
HP, N184, N4H5S2
K.MVSHHN[+2204.772]LTTGATLINEQWLLTTAK.N
HP, N241, N4H5S2
K.VVLHPN[+2204.772]YSQVDIGLIK.L
HP, N207, N4H5S2
K.NLFLN[+2204.772]HSENATAK.D
HP, N184, N4H5S2F1
K.MVSHHN[+2350.830]LTTGATLINEQWLLTTAK.N
HP, N184, N5H6S3
K.MVSHHN[+2861.000]LTTGATLINEQWLLTTAK.N
HPX, N187, N4H5S2
R.SWPAVGN[+2204.772]C[+57.021]SSALR.W
IGHA1, N340, N4H5S2F1
R.LAGKPTHVN[+2350.830]VSVVMAEVDGTC[+57.021].Y
IGHA1, N340, N5H5S1F1
R.LAGKPTHVN[+2262.814]VSVVMAEVDGTC[+57.021].Y
IGHA1, N340, N5H5S2F1
R.LAGKPTHVN[+2553.910]VSVVMAEVDGTC[+57.021].Y
IGHA1;IGHA2, N144, N2H5
R.LSLHRPALEDLLLGSEAN[+1216.423]LTC[+57.021]TLTGLR.D
IGHA1;IGHA2, N144, N4H4S1
R.LSLHRPALEDLLLGSEAN[+1751.624]LTC[+57.021]TLTGLR.D
IGHA1;IGHA2, N144, N5H3
R.LSLHRPALEDLLLGSEAN[+1501.555]LTC[+57.021]TLTGLR.D
IGHA1;IGHA2, N144, N5H5S1
R.LSLHRPALEDLLLGSEAN[+2116.756]LTC[+57.021]TLTGLR.D
ORM1, N56, N4H5S2
K.WFYIASAFRNEEYN[+2204.772]K.S
SERPINA1, N271, N4H5S1
K.YLGN[+1913.677]ATAIFFLPDEGKLQHLENELTHDIITK.F
SERPINA1, N70, N4H5S2
R.QLAHQSN[+2204.772]STNIFFSPVSIATAFAMLSLGTK.A
SERPINA1, N271, N4H5S2
K.YLGN[+2204.772]ATAIFFLPDEGK.L
TF, N630, N4H5S2
R.QQQHLFGSN[+2204.772]VTDC[+57.021]SGNFC[+57.021]LFR.S
HCD Relative Intensity
CID Relative Intensity
m/z

**Figure S4:** Back-to-back comparison of DDA (top panels, HCD, 1.6 m/z window) and DIA (bottom panels, CID, 2 m/z window) MS/MS spectra for each of the candidate glycopeptides from the COVID-19 cohort. For CID/HCD spectra, fragments matched to theoretical fragments exported from Byonic for each DDA spectrum are shown (0.1 Da tolerance). Fragments shared between DDA and DIA spectra are shown in green, oxonium ions in red and singly-assigned fragments in grey.

**Table S1:** IgG glycopeptides identified by OxoScan-MS analysis.

| IgG | Glycan | m/z | Monoisotopic Mass | Charge | RT | [M+H]+ | peak_num |
|---|---|---|---|---|---|---|---|
| 1 | G0 | 830.0054 | 2486.9943 | 3 | 27.55 | 2488.0016 | 17 |
| 1 | G0F | 878.6832 | 2633.0277 | 3 | 26.85 | 2634.035 | 7 |
| 1 | G0FN | 946.3678 | 2836.0815 | 3 | 27.95 | 2837.0888 | 14 |
| 1 | G1F | 932.6988 | 2795.0745 | 3 | 26.55 | 2796.0818 | 4 |
| 1 | G1F | 932.7058 | 2795.0955 | 3 | 26.96 | 2796.1028 | 18 |
| 1 | G1FN | 1000.3946 | 2998.1619 | 3 | 27.75 | 2999.1692 | 13 |
| 1 | G1FS1 | 1029.7458 | 3086.2155 | 3 | 28.75 | 3087.2228 | 21 |
| 1 | G2F | 986.7129 | 2957.1168 | 3 | 26.46 | 2958.1241 | 8 |
| 1 | G2FN | 1054.4152 | 3160.2237 | 3 | 27.55 | 3161.231 | 20 |
| 1 | G2FS1 | 1083.7589 | 3248.2548 | 3 | 28.15 | 3249.2621 | 11 |
| 2 | G0 | 819.3432 | 2455.0077 | 3 | 52.15 | 2456.015 | 35 |
| 2 | G0F | 868.0212 | 2601.0417 | 3 | 50.95 | 2602.049 | 1 |
| 2 | G0FN | 935.703 | 2804.0871 | 3 | 51.25 | 2805.0944 | 9 |
| 2 | G1F | 922.0331 | 2763.0774 | 3 | 50.15 | 2764.0847 | 3 |
| 2 | G1F | 922.031 | 2763.0711 | 3 | 50.75 | 2764.0784 | 5 |
| 2 | G1FN | 989.7264 | 2966.1573 | 3 | 50.55 | 2967.1646 | 12 |
| 2 | G1FN | 989.7217 | 2966.1432 | 3 | 50.95 | 2967.1505 | 27 |
| 2 | G1FS1 | 1019.0768 | 3054.2085 | 3 | 53.65 | 3055.2158 | 10 |
| 2 | G2F | 976.0541 | 2925.1404 | 3 | 49.85 | 2926.1477 | 2 |
| 2 | G2FN | 1043.7621 | 3128.2644 | 3 | 50.25 | 3129.2717 | 15 |
| 2 | G2FS1 | 1073.0854 | 3216.2343 | 3 | 52.75 | 3217.2416 | 6 |
| 4 | G0F | 873.36 | 2617.0581 | 3 | 38.05 | 2618.0654 | 19 |
| 4 | G0FN | 941.0292 | 2820.0657 | 3 | 38.55 | 2821.073 | 33 |
| 4 | G1F | 927.3766 | 2779.1079 | 3 | 37.45 | 2780.1152 | 23 |
| 4 | G1F | 927.3892 | 2779.1457 | 3 | 37.15 | 2780.153 | 69 |
| 4 | G1F | 927.3757 | 2779.1052 | 3 | 37.95 | 2780.1125 | 43 |
| 4 | G1FN | 995.058 | 2982.1521 | 3 | 38.15 | 2983.1594 | 36 |
| 4 | G2F | 981.38 | 2941.1181 | 3 | 37.24 | 2942.1254 | 26 |
| 4 | G2FN | 1049.0927 | 3144.2562 | 3 | 37.85 | 3145.2635 | 81 |
| 4 | G2FS1 | 1078.4419 | 3232.3038 | 3 | 39.55 | 3233.3111 | 24 |

**Table S2:** COVID-19 clinical cohort demographics.

| Sample ID | Age | Sex | WHO Grade | Clinical Category | Sample ID | Age | Sex | WHO Grade | Clinical Category | Sample ID | Age | Sex | WHO Grade | Clinical Category |
|---|---|---|---|---|---|---|---|---|---|---|---|---|---|---|
| HD-1 | 27 | w | 0 | Healthy | CV-1 | 22 | m | 3 | Mild | CV-16 | 74 | m | 5 | Moderate |
| HD-2 | 23 | m | 0 | Healthy | CV-19 | 52 | m | 3 | Mild | CV-63 | 61 | m | 5 | Moderate |
| HD-3 | 34 | w | 0 | Healthy | CV-23 | 44 | w | 3 | Mild | CV-8 | 63 | m | 7 | Severe |
| HD-4 | 31 | w | 0 | Healthy | CV-35 | 37 | w | 3 | Mild | CV-9 | 80 | w | 7 | Severe |
| HD-5 | 30 | m | 0 | Healthy | CV-37 | 78 | m | 3 | Mild | CV-13 | 71 | m | 7 | Severe |
| HD-6 | 41 | w | 0 | Healthy | CV-49 | 72 | w | 3 | Mild | CV-25 | 45 | w | 7 | Severe |
| HD-7 | 24 | w | 0 | Healthy | CV-56 | 58 | w | 3 | Mild | CV-32 | 50 | w | 7 | Severe |
| HD-8 | 27 | m | 0 | Healthy | CV-65 | 22 | m | 3 | Mild | CV-33 | 35 | w | 7 | Severe |
| HD-9 | 30 | w | 0 | Healthy | CV-66 | 75 | w | 3 | Mild | CV-57 | 55 | w | 6 | Severe |
| HD-10 | 34 | w | 0 | Healthy | CV-75 | 84 | m | 3 | Mild | CV-58 | 26 | m | 7 | Severe |
| HD-11 | 42 | m | 0 | Healthy | CV-11 | 61 | m | 5 | Moderate | CV-59 | 86 | m | 7 | Severe |
| HD-12 | 30 | w | 0 | Healthy | CV-24 | 64 | m | 4 | Moderate | CV-60 | 65 | m | 6 | Severe |
| HD-13 | 33 | w | 0 | Healthy | CV-38 | 70 | m | 4 | Moderate | CV-61 | 54 | m | 7 | Severe |
| HD-14 | 29 | w | 0 | Healthy | CV-48 | 48 | m | 4 | Moderate | CV-62 | 72 | m | 6 | Severe |
| HD-15 | 36 | w | 0 | Healthy | CV-50 | 78 | m | 4 | Moderate | CV-64 | 57 | m | 7 | Severe |

**Table S3:** Non-linear chromatographic gradient for COVID-19 cohort method.

| Time | Flow rate (µl/min) | %A | %B |
|---|---|---|---|
| 0.00 | 5 | 97.0 | 3.0 |
| 0.86 | 5 | 92.9 | 7.1 |
| 2.42 | 5 | 88.8 | 11.2 |
| 5.53 | 5 | 84.7 | 15.3 |
| 9.38 | 5 | 80.6 | 19.4 |
| 13.02 | 5 | 76.4 | 23.6 |
| 15.48 | 5 | 72.3 | 27.7 |
| 17.27 | 5 | 68.2 | 31.8 |
| 19.00 | 5 | 60.0 | 40.0 |
| 20.00 | 5 | 20.0 | 80.0 |
| 20.50 | 5 | 20.0 | 80.0 |
| 21.50 | 5 | 97.0 | 3.0 |
| 27.50 | 5 | 97.0 | 3.0 |

**Table S4:** Scanning SWATH parameters for different gradients. All chromatographic gradients are linear (3-40% Buffer B) except that described in Table S2.

| Method Name | Active Gradient Length (min) | Precursor Range (m/z) | Window Width (m/z) | Cycle Time (s) | Effective Accumulation Time (ms) |
|---|---|---|---|---|---|
| Cohort | 19 | 800-1400 | 10 | 1.5 | 21.7 |
| Deglyco | 45 | 1000-1500 | 5 | 3 | 28.2 |
| IgG | 90 | 800-1600 | 10 | 6 | 71.5 |
| GPF-1 | 90 | 500-1000 | 1 | 6 | 11.7 |
| GPF-2 | 90 | 1000-1500 | 1 | 6 | 11.7 |
| GPF-3 | 90 | 1500-2000 | 1 | 6 | 11.7 |

**Table S5:** Glycopeptides identified in DDA experiments and quantified across the COVID-19 cohort by OxoScan-MS.

| m/z | Charge | Gene | Peptide | Glycan | Position | Monoisotopic Mass | peak_num |
|---|---|---|---|---|---|---|---|
| 1326.2433 | 3 | AHSG | K.VC[+57.021]QDC[+57.021]PLLAPLN[+2204.772]DTR.V | N4H5S2 | N156 | 3975.708 | 98 |
| 891.4581 | 3 | AHSG | R.TVVQPS[+656.228]VGAAAGPVVPPC[+57.021]PGR.I | N1H1S1 | S346 | 2671.3524 | 66 |
| 1269.5496 | 3 | FGB | R.GTAGNALMDGASQLMGEN[+1913.677]R.T | N4H5S1 | N394 | 3805.6269 | 8 |
| 1366.556 | 3 | FGB | R.GTAGNALMDGASQLMGEN[+2204.772]R.T | N4H5S2 | N394 | 4096.6461 | 11 |
| 1172.5169 | 3 | FGB | R.GTAGNALMDGASQLMGEN[+1622.582]R.T | N4H5 | N394 | 3514.5288 | 203 |
| 1009.9792 | 4 | FGG | K.VDKDLQSLEDILHQVEN[+1913.677]K.T | N4H5S1 | N78 | 4035.8876 | 20 |
| 937.2088 | 4 | FGG | K.VDKDLQSLEDILHQVEN[+1622.582]K.T | N4H5 | N78 | 3744.806 | 499 |
| 1221.8325 | 4 | HP | K.MVSHHN[+2204.772]LTTGATLINEQWLLTTAK.N | N4H5S2 | N184 | 4883.3008 | 1 |
| 1385.8938 | 4 | HP | K.MVSHHN[+2861.000]LTTGATLINEQWLLTTAK.N | N5H6S3 | N184 | 5539.546 | 23 |
| 1258.352 | 4 | HP | K.MVSHHN[+2350.830]LTTGATLINEQWLLTTAK.N | N4H5S2F1 | N184 | 5029.3788 | 125 |
| 1221.8687 | 3 | HP | K.NLFLN[+2204.772]HSENATAK.D | N4H5S2 | N207 | 3662.5842 | 292 |
| 1333.9684 | 3 | HP | K.VVLHPN[+2204.772]YSQVDIGLIK.L | N4H5S2 | N241 | 3998.8833 | 3 |
| 1203.8456 | 3 | HPX | R.SWPAVGN[+2204.772]C[+57.021]SSALR.W | N4H5S2 | N187 | 3608.5149 | 19 |
| 1134.5156 | 4 | IGHA1 | R.LAGKPTHVN[+2350.830]VSVVMAEVDGTC[+57.021].Y | N4H5S2F1 | N340 | 4534.0332 | 73 |
| 1185.275 | 4 | IGHA1 | R.LAGKPTHVN[+2553.910]VSVVMAEVDGTC[+57.021].Y | N5H5S2F1 | N340 | 4737.0708 | 50 |
| 1112.5078 | 4 | IGHA1 | R.LAGKPTHVN[+2262.814]VSVVMAEVDGTC[+57.021].Y | N5H5S1F1 | N340 | 4446.002 | 172 |
| 1179.6014 | 4 | IGHA1;IGHA2 | R.LSLHRPALEDLLLGSEAN[+1751.624]LTC[+57.021]TLTGLR.D | N4H4S1 | N144 | 4714.3764 | 40 |
| 1270.8887 | 4 | IGHA1;IGHA2 | R.LSLHRPALEDLLLGSEAN[+2116.756]LTC[+57.021]TLTGLR.D | N5H5S1 | N144 | 5079.5256 | 16 |
| 1045.7969 | 4 | IGHA1;IGHA2 | R.LSLHRPALEDLLLGSEAN[+1216.423]LTC[+57.021]TLTGLR.D | N2H5 | N144 | 4179.1584 | 130 |
| 1117.0811 | 4 | IGHA1;IGHA2 | R.LSLHRPALEDLLLGSEAN[+1501.555]LTC[+57.021]TLTGLR.D | N5H3 | N144 | 4464.2952 | 28 |
| 1028.8025 | 3 | IGHG2 | K.TKPREEQFN[+1444.534]STFR.V | N4H3F1 | N176 | 3083.3856 | 35 |
| 1036.459 | 4 | ORM1 | K.WFYIASAFRNEEYN[+2204.772]K.S | N4H5S2 | N56 | 4141.8068 | 142 |
| 1320.9347 | 3 | SERPINA1 | K.YLGN[+2204.772]ATAIFFLPDEGK.L | N4H5S2 | N271 | 3959.7822 | 26 |
| 1091.7508 | 5 | SERPINA1 | K.YLGN[+1913.677]ATAIFFLPDEGKLQHLENELTHDIITK.F | N4H5S1 | N271 | 5453.7175 | 39 |
| 1347.4086 | 4 | SERPINA1 | R.QLAHQSN[+2204.772]STNIFFSPVSIATAFAMLSLGTK.A | N4H5S2 | N70 | 5385.6052 | 5 |
| 1180.7626 | 4 | TF | R.QQQHLFGSN[+2204.772]VTDC[+57.021]SGNFC[+57.021]LFR.S | N4H5S2 | N630 | 4719.0212 | 12 |
